# A drug-tunable gene therapy for broad-spectrum protection against retinal degeneration

**DOI:** 10.1101/283887

**Authors:** Clayton P. Santiago, Casey J. Keuthan, Sanford L. Boye, Shannon E. Boye, Aisha A. Imam, John D. Ash

## Abstract

Retinal degenerations are a large cluster of diseases characterized by the irreversible loss of light-sensitive photoreceptors that impairs the vision of 9.1 million people in the US. An attractive treatment option is to use gene therapy to deliver broad-spectrum neuroprotective factors. However, this approach has had limited clinical translation because of the inability to control transgene expression. To address this problem, we generated an adeno-associated virus vector named RPF2 that was engineered to express domains of leukemia inhibitory factor fused to the destabilization domain of bacterial dihydrofolate reductase. Fusion proteins containing the destabilization domain are degraded in mammalian cells but can be stabilized with the binding of the drug trimethoprim. Our data show that expression levels of RPF2 are tightly regulated by the dose of trimethoprim, and can be reversed by trimethoprim withdrawal. We further show that stabilized RPF2 can protect photoreceptors and prevent blindness in treated mice.

## Introduction

Vision loss is often due to the death of neurons that either initiate the visual response to light (photoreceptors), or the neurons that transmit the signal to the brain (bipolar cells, ganglion cells, etc) (1). As there are approximately 18 million people worldwide living with some form of retinal degenerating disease, this is a significant cause of morbidity in human health (2). To date, mutations in approximately 250 genes have been identified to cause inherited retinal diseases (3). For example, retinitis pigmentosa is a group of monogenetic diseases in which patients often have impaired night and peripheral vision beginning from childhood, which progressively worsen until central vision is lost (4). Other retinal degenerative disorders such as age-related macular degeneration (AMD) are caused by a multitude of factors such as polymorphisms in complement factor H, obesity, smoking and hypertension (5–10). To address these diseases, many gene therapy strategies, including targeted gene replacement and gene editing, are under development (11–15). However, it is a daunting task to design therapies for every individual gene or mutation that causes retinal dysfunction. To circumvent this challenge, additional strategies are based on gene/mutation-independent approaches, including expression of transcriptional regulators, inhibitors of apoptosis, regulators of oxidative defense, inhibitors of inflammation, and expression of neurotrophic factors (16–24).

We, and others, have found that neuroprotective cytokines such as leukemia inhibitory factor (LIF) or ciliary neurotrophic factor (CNTF) can protect photoreceptors from a broad range of insults, including mechanical injury and multiple mutations that cause inherited retinal degeneration (25–27). However, one problem with using LIF or CNTF as a gene therapy is that the long-term effects of highly expressed neurotrophic factors have been shown to be detrimental to retinal function and may promote inflammation (28–30). What is needed to make these gene-independent approaches a viable therapy is a mechanism to control the level of transgene expression and the ability to turn it off, should adverse events occur.

Recently, drug-regulated destabilization domains have been shown to control transgene expression in the brain (31–33). This system requires fusing the *E. coli* dihydrofolate reductase (DHFR) gene to the transgene of interest to produce a fusion protein. In mammalian cells, fusion proteins containing the DHFR protein are rapidly ubiquitinated and degraded by the proteasome system (34, 35). Because the DHFR portion of the fusion protein causes protein turnover, it is referred to as a destabilization domain (DD). The FDA-approved antibiotic, Trimethoprim (TMP), can bind to the DD and prevent the protein from being degraded, which allows the fusion protein to escape degradation (31). Additionally, this antibiotic can cross the blood-brain barrier (32, 33).

In this study, we designed a neurotrophic factor-DD fusion protein that we have named Retinal Protective Factor 2 (RPF2). Our data demonstrate that TMP can be used to regulate RPF2 production *in vitro* and *in vivo* in a dose-dependent manner and that expression was reversible by TMP withdrawal. Moreover, we found that long-term treatment of RPF2 did not alter retinal function or morphology. Lastly, RPF2 treatment could rescue photoreceptors from an acute light-induced degeneration model, as well as preserve cone vision in the phosphodiesterase 6b mutant (*rd10*) inherited retinal degeneration model.

## Results

### Unregulated, long-term cytokine expression is detrimental to the retina

In this study, we utilized an AAV2-mutant capsid to deliver the human LIF (hLIF) transgene to the retina by intravitreal injection. This capsid contains four proteosomal avoidance mutations and is referred to as AAV2 (quadY-F+T-V). AAV2 (quadY-F+T-V) was selected since it can efficiently transduce a broad range of cell types in the murine retina by intravitreal injection (Supplemental Figure 1A) (36). Additionally, the AAV2 capsid is the only AAV serotype that can effectively transduce retinal cells in non-human primates (37) .

An AAV2 (quadY-F+T-V) vector expressing the hLIF transgene under the control of the chicken beta-actin (CBA) promoter was injected into the vitreous of Balb/cJ mice at two titers (1×10^8^ vector genomes/µL for low titer, 1×10^9^ vector genomes/µL for high titer) (Figure 1A). After seven weeks, using a well-established light damage (LD) model, mice were exposed to damaging bright light that was calibrated to synchronously kill the majority of photoreceptors. Mice injected with AAV2 (quadY-F+T-V)-hLIF had significant preservation of photoreceptors in the outer nuclear (ONL) compared to controls (Supplemental Figure 2). However, despite this protective activity, long-term studies on treated mice not exposed to damaging light had progressive, abnormal retinal structures consistent with edema, as seen by optical coherence tomography (OCT) imaging (Figure 1B-D). Long-term expression also caused a reduction in photoreceptor light responses, as measured by electroretinography (ERG) (Figure 1E and F). We observed edema in animals treated with the low titer of AAV2 (quadY-F+T-V)-hLIF, which was significantly worse in the high titer group (Supplemental Figure 3A). The severity of adverse retinal changes (edema and loss of function) correlated with the level of LIF protein expression and activation of STAT3 (Supplemental Figure 3B and C). Overall, these results demonstrate that unregulated, high levels of LIF expression would be of limited therapeutic value.

**Figure 1.**
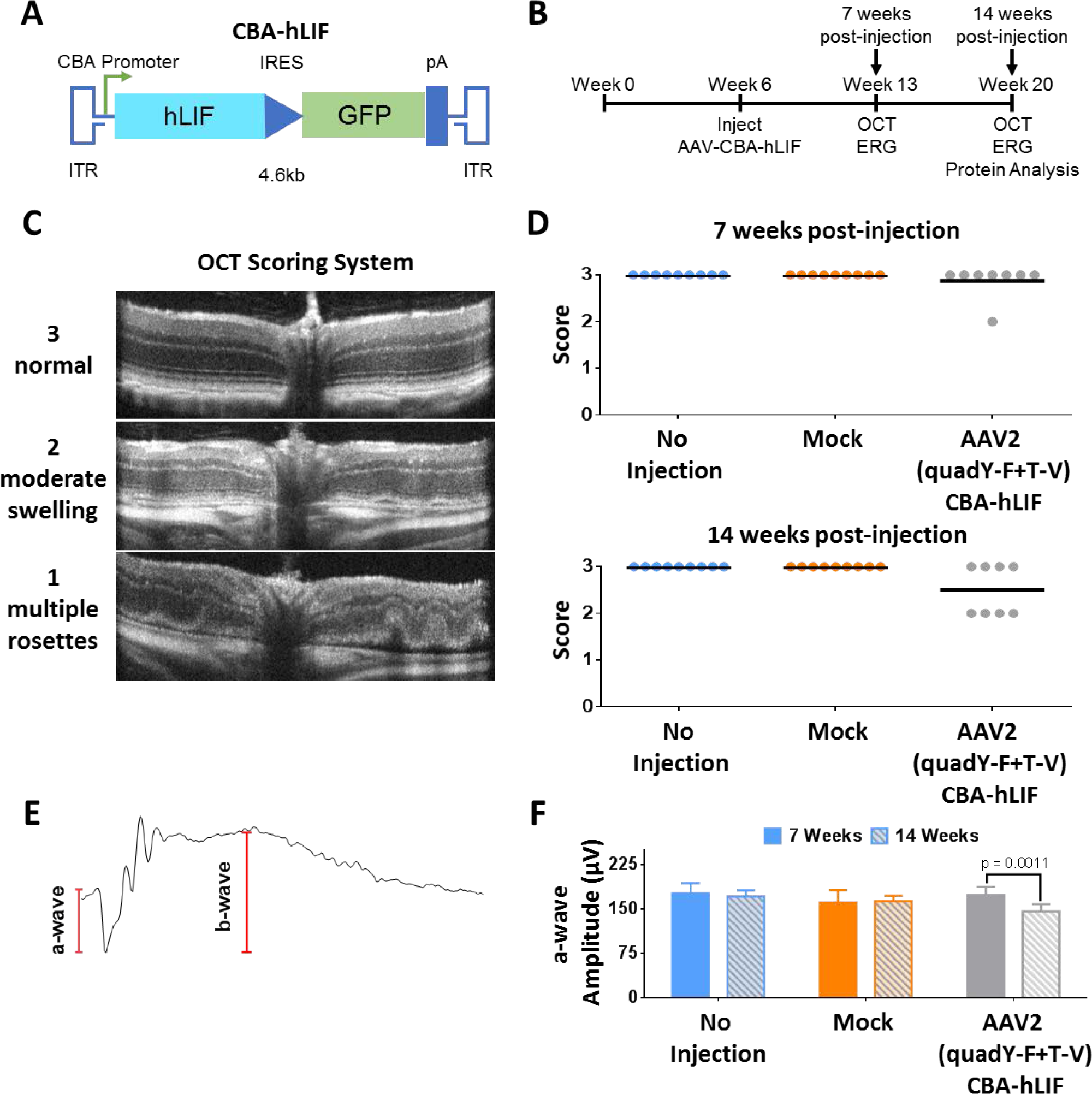
Long-term constitutive expression of AAV-hLIF leads can have adverse effects on retinal morphology and function. (A) Schematic of AAV-hLIF construct showing the human LIF (hLIF) transgene under the control of the chicken beta actin (CBA) promoter. The expression construct is flanked by inverted terminal repeats (ITRs) to allow for packaging into AAV vectors. The size of the vector is 4.6kb. **(B)** Experimental timeline using the AAV-hLIF vector. At 6 weeks of age, Balb/c mice were intravitreally injected with AAV2 (quadY-F+T-V) carrying the hLIF transgene. *In vivo* imaging and functional analysis was performed 7 weeks and 14 weeks post-injection. **(C)** Scoring rubric for assessment of retinal morphology. A score of 3 indicated a normal retina that maintains well-defined boundaries of hyper or hypo reflectance. A score of 2 indicated retinas with minor ruffling and less-defined retinal layer boundaries. Retinas with a score of 1 had multiple rosettes, retinal thinning, excessive cells in the vitreous, and retinal detachment. **(D)** Retinal scoring in control and AAV2 (quadY-F+T-V) hLIF-injected animals 7 (top) and 14 (bottom) weeks post-injection. Each filled circle represents one animal (n = 8-9) and the black lines represent the mean for each group. **(E)** Representative ERG trace. The a-wave, the initial negative deflection, reflects the hyperpolarization of the photoreceptors following light stimulation. The b-wave amplitude was measured from the trough of the a wave to the adjacent peak, indicative of the depolarization due to voltage changes taking place in the inner retina. **(F)** Average a-wave amplitudes at 1.0 cd.s/m^2^ of control and AAV2 (quadY-F+T-V) hLIF-injected animals 7 and 14 weeks post-injection. Error bars represent SEM for each group. One-way ANOVA with Sidak post-hoc test was performed; n = 8 animals per group.

### Development of Retinal Protective Factor 2

In our initial attempt to regulate LIF expression, a synthetic and codon-optimized *E. coli* DHFR (DD) cDNA was fused to the hLIF cDNA (Supplemental Figure 4A). While the resulting fusion protein was expressed in cell culture, it was not regulated by TMP and was unable to activate STAT3 downstream of its receptor (Supplemental Figure 4B and C). Our interpretation of these results was that both domains in the fusion protein inhibited the activity of the other. In attempts to produce a regulatable and functional cytokine, we added either 1) sequences encoding an eight amino acid linker with one furin cleavage site (FCS) between hLIF and the DD, or 2) sequences encoding a twelve amino acid linker with 2xFCS (Figure 2A and Supplemental Figure 4A). This strategy takes advantage of the furin protease system that cleaves the domains as they transit the secretory pathway (38). The fusion proteins were tested *in vitro*, and the 12 amino acid linker containing 2xFCS produced the highest levels of cleaved cytokine with TMP treatment and had minimally detectable levels of cytokine in the absence of TMP (Figure 2C). In addition, the expression of cleaved and secreted cytokine was regulated by TMP in a dose-dependent manner (Supplemental Figure 4D). The effects of TMP were dependent on the DD fusion, since TMP did not alter the expression of cytokine in the psiCHECK or hLIF-transfected group. To determine whether the secreted RPF2 was functional, the conditioned media from transfected cells were collected and used to stimulate STAT3 in untransfected rat Müller glial-like (rMC-1) cells. Only conditioned media from TMP-treated cells led to a significant increase in STAT3 activation (Figure 2D). These levels were comparable to that of the hLIF-transduced group. Our *in vitro* data suggest that RPF2 can be regulated in a dose-dependent manner and can be secreted in its functional form to activate STAT3. Because of its superior regulation the fusion protein with 2xFCS, named Retinal Protective Factor 2 (RPF2) was packaged into AAV for further testing.

**Figure 2.**
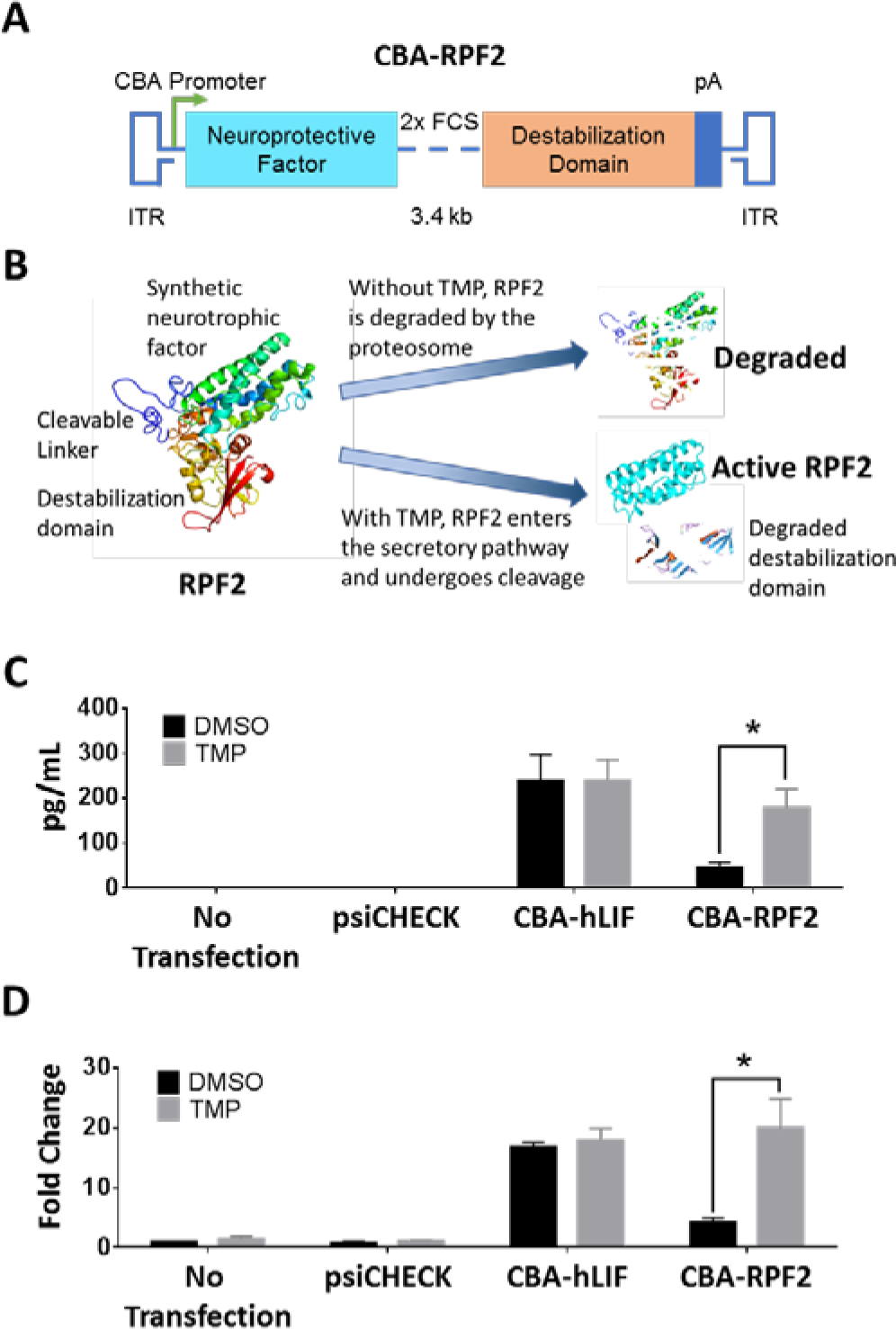
TMP-stabilized RPF2 is active *in vitro.* (A) Schematic of the RPF2 construct showing the synthetic neuroprotective factor linked to the destabilization domain expressed from the CBA promoter. Within the linker region, two furin cleavage sites (FCS) separate the domains. **(B)** Illustration of the RPF2 stabilization strategy. In the absence of TMP, RPF2 is rapidly degraded by the proteasome following translation. TMP stabilizes RPF2, allowing it to be processed and secreted. **(C)** ELISA measuring cytokine levels in the media of rMC-1 cells transfected with CBA-RPF2 with or without TMP. **(D)** Immunoblot analysis of pSTAT3 Y705 levels in rMC-1 cells following treatment with conditioned media from CBA-RPF2-transfected cells, with and without TMP treatment. Error bars represent SEM (C-D). Two-way ANOVA with Sidak post-hoc test was performed for C-D; n = 4 biological replicates; * p < 0.0001.

### AAV-RPF2 is regulated by TMP in a dose-dependent manner *in vivo*

RPF2 was packaged into the AAV2 (quadY-F+T-V) capsid and intravitreally injected into Balb/cJ mice at high and low titers. Six weeks after AAV injections, mice were given seven daily IP injections of varying doses of TMP. RPF2 levels rose with increasing doses of TMP and fell to baseline following a 7-day TMP withdrawal (Figure 3A). Cytokine levels for both the TMP-vehicle only (0 mg/kg) and TMP withdrawal groups were below the limit of detection for the assay. The expressed cytokine was also functional since STAT3 activation in the retina was higher in animals treated with TMP (data not shown). To characterize the long-term effects of the AAV-mediated RPF2 expression, retinal structure and function were monitored over time following intravitreal injection (Figure 3B). Animals were placed on chow containing TMP starting at 7 weeks post-injection. This route of drug administration yielded RPF2 stabilization at levels comparable to that of the 60 mg/kg systemic TMP delivery and did not affect the animal’s daily food intake (Supplemental Figure 5B and 6). After 7 weeks of TMP treatment, mice injected with AAV2 (quadY-F+T-V)-RPF2 had normal retinal morphology and clearly lacked the edema observed in the AAV2 (quadY-F+T-V)-hLIF treated mice (Figure 3C and Supplemental Figure 5C). Mice also had normal photoreceptor function, as reflected in scotopic ERG responses (Figure 3D). These data show that AAV-mediated RPF2 expression can be regulated by TMP in a dose-dependent manner *in vivo*, and long-term expression did not result in the negative outcomes that we observed using AAV2 (quadY-F+T-V)-hLIF.

**Figure 3.**
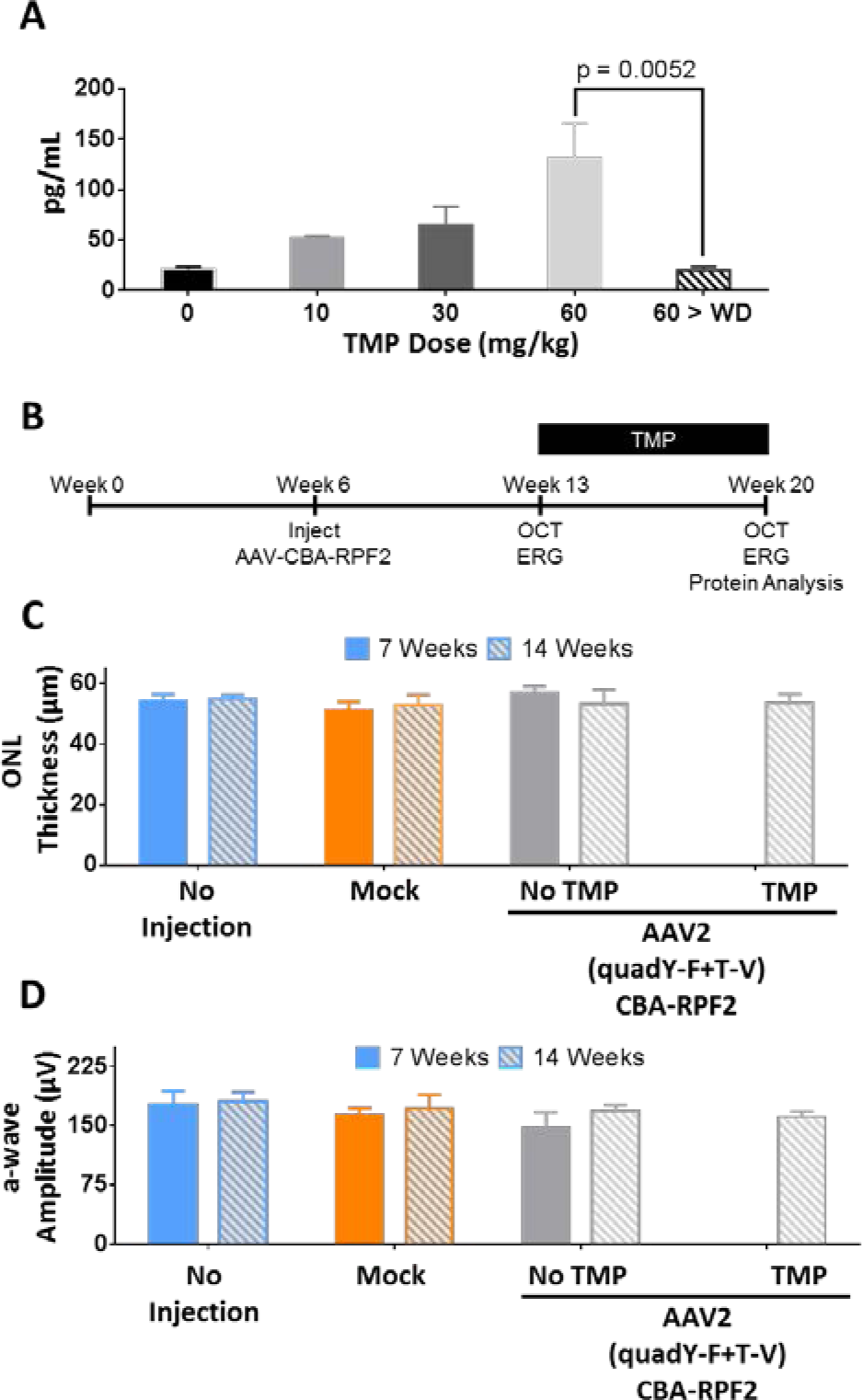
RPF2 is regulated in a dose-dependent manner and demonstrates no toxicity *in vivo*. (A) ELISA levels of RPF2 in the retinal lysates of animals injected with low titer AAV2 (quadY-F+T-V) RPF2 at varying concentrations of TMP. Tissue was also collected 7 days after TMP withdrawal (60 > WD). One-way ANOVA with Sidak post-hoc test was performed; n = 4 biological replicates. **(B)** Experimental timeline using the AAV2 (quadY-F+T-V) RPF2 vectors following a similar strategy to that in Figure 1B. From Week 13 to Week 20, mice were kept on either normal chow or a TMP-supplemented diet. **(C)** Average outer nuclear layer (ONL) thickness in injected animals before and after TMP diet. **(D)** Average a-wave amplitudes at 1.0 cd.s/m^2^ of control and AAV2 (quadY-F+T-V) RPF2-injected animals 7 and 14 weeks post-injection. Two-way ANOVA with Sidak post-hoc test was performed for C-D; n = 8 biological replicates. No statistical significance was found.

### AAV-RPF2 can protect photoreceptors from acute light injury

To determine whether an AAV-mediated RPF2 therapy could protect photoreceptors from LD, six-week-old mice were injected with AAV2 (quadY-F+T-V)-RPF2 at low and high titers. After 7 weeks, mice were given TMP in mouse chow for 7 days prior to exposure to damaging bright light (Figure 4A). In the presence of TMP, animals injected with AAV2 (quadY-F+T-V)-RPF2 showed significant preservation of ONL thickness compared to mock-injected or no TMP control groups, suggesting that the TMP-induced cytokine protected photoreceptors (Figure 4B). TMP-induced RPF2 not only preserved structure, but also preserved photoreceptor function (Figure 4C). High-titer AAV-RPF2 groups had greater preservation of the ONL than the low titer groups (Supplemental Figure 7A-C). Overall, the AAV-RPF2 therapy successfully protected the murine retinas from acute light injury.

**Figure 4.**
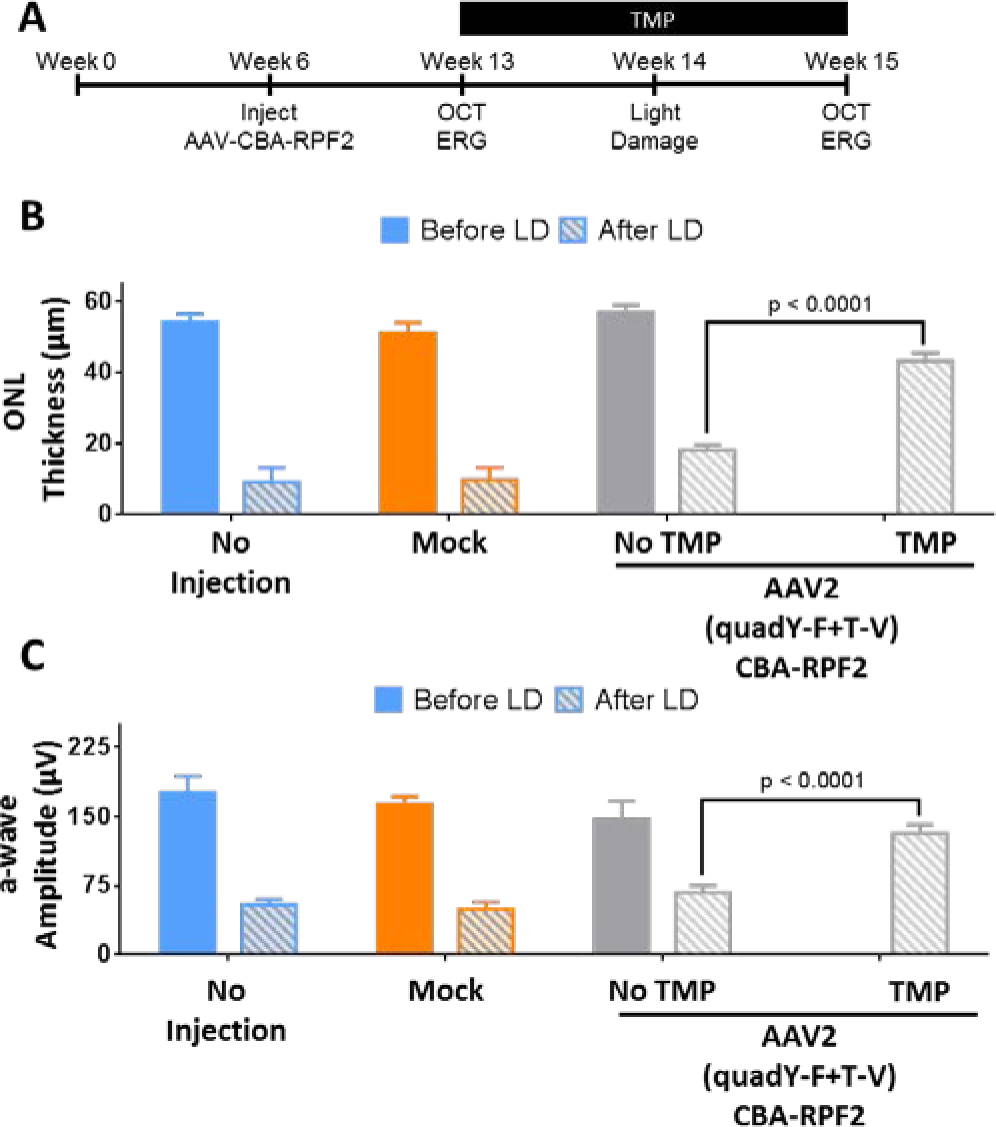
RPF2 can reduce retinal degeneration in an acute light damage model. (A) Experimental timeline using the AAV2 (quadY-F+T-V) RPF2 vectors with light damage (LD). One week after the start of TMP supplementation, animals were LD-treated, followed by a week of recovery to allow of clearance of damaged cells. **(B)** Average ONL thickness in injected animals before and after LD treatment. **(C)** Average a-wave amplitudes at 1.0 cd.s/m^2^ of control and AAV-RPF2-injected animals before and after LD treatment. Error bars indicate SEM for B-C. Two-way ANOVA with Sidak post-hoc test was performed for B-C; n = 8-9 biological replicates per group.

### AAV-RPF2 protects photoreceptors in a model of inherited retinal degeneration

The *rd10* mouse is a widely used model to study inherited retinal degeneration (39–43). To determine whether TMP-induced RPF2 expression could preserve photoreceptors in *rd10* retinas, AAV2 (quadY-F+T-V)-RPF2 was intravitreally injected at high titer to the animals at 2 weeks of age, prior to any degeneration (Figure 5A). At week 3 (weaning age), TMP was administered and the animals were monitored by OCT and ERG over time. The TMP-treated AAV-RPF2 groups had significant preservation of the ONL compared to no TMP controls (Figure 5B). Retinas were collected at week 8 to measure preservation of cones, which normally die subsequent to rod loss. In the AAV-RPF2 groups treated with TMP, cones were markedly preserved compared to mock-injected mice or no TMP controls (Figure 5C and D). The preserved cones maintained majority of their function in the TMP-treated AAV-RPF2 groups, as measured by photopic ERG responses (Figure 6A). To quantify functional vision in treated mice, we used OptoMotry to measure optokinetic reflex (Figure 6B) (44, 45). The AAV2 (quadY-F+T-V)-RPF2 group treated with TMP showed significant preservation of cone-mediated visual behavior compared to mock-injected or no-TMP groups. Responses in TMP-treated mice were comparable to that of C57BL/6J wild-type control mice (Figure 6C). Overall, our data show AAV-mediated RPF2 expression is capable of protecting both rods and cones and preserving cone vision in this aggressive model of inherited retinal degeneration.

**Figure 5.**
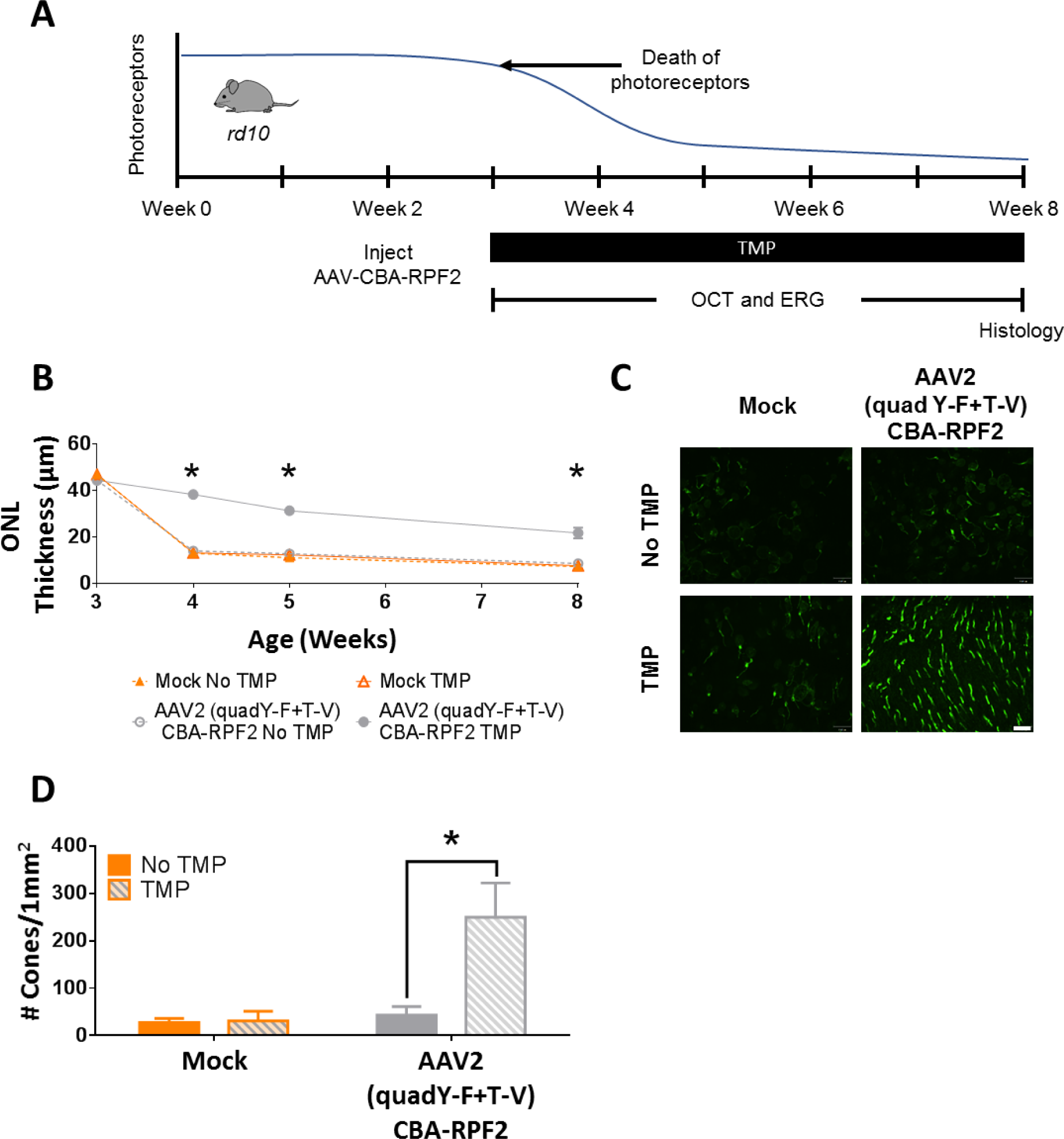
RPF2 can preserve cone morphology in the inherited *rd10* retinal degeneration model. (A) Timeline showing the natural history of the *rd10* mouse model and therapeutic intervention with AAV2 (quadY-F+T-V) RPF2 vectors. The onset of photoreceptor degeneration begins at Week 3. Animals were injected with AAV2 (quadY-F+T-V) RPF2 vectors at 2 weeks of age. At weaning, animals were placed on a TMP diet. Morphological and functional analysis was performed over the subsequent weeks before collecting the eyes for histology. **(B)** Average ONL thickness in therapy-treated animals over time. Two-way ANOVA with Sidak post-hoc test was performed for B and D; n = 6 biological replicates per group; * p < 0.0001. Error bars indicate SEM. **(C)** Immunofluorescent images of retinal flat mounts showing staining of s and m opsin (green). Images were taken at 40X magnification. White bar indicates 100µm. **(D)** Quantification of cones in C. Error bars indicate SEM. Two-way ANOVA with Sidak post-hoc test was performed for B and D; n = 4 biological replicates per group; * p < 0.0001.

**Figure 6.**
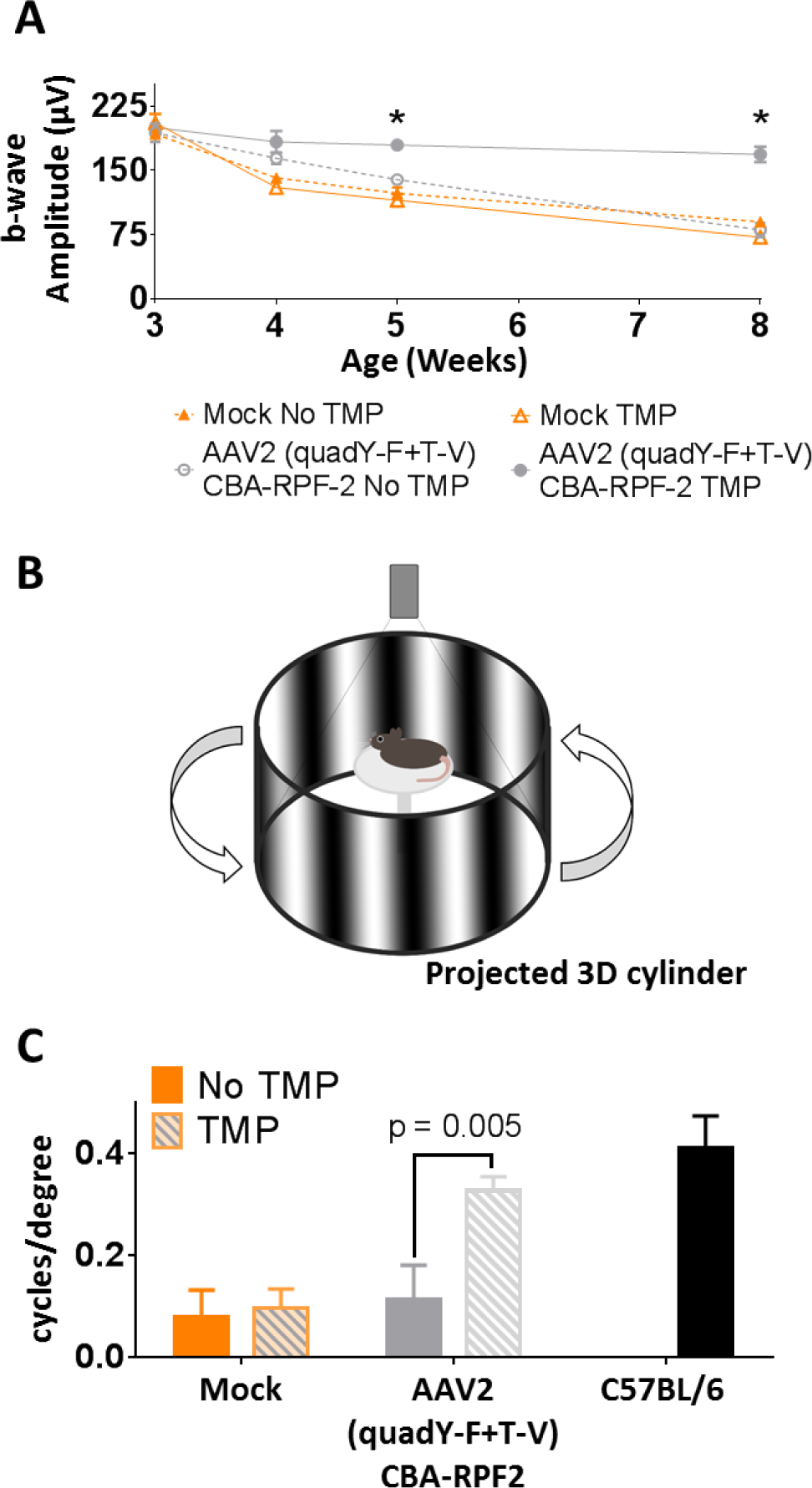
RPF2-treated *rd10* mice maintain long term cone-dependent vision. (A) Average b-wave amplitudes at photopic 60 cd.s/m^2^ of control and AAV2 (quadY-F+T-V) RPF2-injected animals over time. **(B)** Illustration of the OptoMotry system used to measure optokinetic tracking response in therapy-treated animals. Animals stand on an elevated platform surrounded by computer monitors that display a sine wave grating, creating a virtual cylinder. The animals track the rotation of the cylinder with reflexive head and neck movements, which is detected by an overhead camera. **(C)** Quantification of optokinetic from OptoMotry analysis described in B. Error bars indicate SEM for A and C. Two-way ANOVA with Sidak post-hoc test was performed for A and C; For A, n = 6 biological replicates per group; * p < 0.0001. For C, n = 4 biological replicates per group except for C57BL/6 = 2 biological replicates.

## Discussion

In this study, we tested RPF2 in two different models of degeneration to determine its ability to protect from a broad-spectrum of insults. The LD model is a well-established method to synchronously induce photoreceptor death (27). Cellular damage is believed to be caused by the formation of free radicals, resulting in cell loss (46). In diseases like AMD, where oxidative stress is believed to be a contributing factor in disease progression, administration of RPF2 could be beneficial. For our second model, we used a mouse with an inherited mutation in the beta subunit of rod phosphodiesterase, a critical component of the phototransduction pathway. This mutation leads to aggressive retinal degeneration, with extensive retinal remodeling within the first month of life. For the maintenance of useful vision in humans, it is of paramount importance to keep cones functional, since humans predominantly utilize cones in their central vision. We showed that AAV-mediated RPF2 expression preserves both rod and cones, and promotes maintenance of useful vision.

Currently, few systems exist to regulate transgene expression delivered by AAV. The most frequently used approach is to limit expression to a particular subset of cells by altering the promoter. Exogenous effector molecules like tetracycline have also been used to induce expression (47). While tetracycline promoters have a wide dynamic range, this system can induce unwanted immune responses in non-human primates (48). Autoregulatory promoters are an alternative to the exogenous effectors (49, 50). However, these cannot be controlled at will, and if the promoters do not express at high levels they are at risk of being silenced over time (51). Additionally, autoregulatory systems are not tunable to meet the personalized needs of the patient and are not reversible. Post-transcriptional control using microRNAs have the same limitations and may have significant off-target effects (52). The TMP-protein regulation approach does not suffer from these weaknesses. Thus far, studies using TMP regulation in the rodent brain have shown that there are no immunogenic properties associated with the destabilization domain (32). Furthermore, this system has the ability to be finely tuned for tight control of transgene expression by adjusting TMP dosing and expression can be reversed if needed, addressing a major barrier in the development of neuroprotective gene therapies. Post-translational regulation has also been shown to be more rapid compared to transcriptional control (53). Combining this approach with a high-expressing promoter, we can reduce the potential silencing effects seen in low-expressing promoters. This makes the system attractive for use in the central nervous system (32, 33).

The use of TMP as a drug to regulate expression also has several advantages. TMP is a FDA-approved drug that is orally bioavailable and can cross the blood-retinal barrier. The TMP dose required to induce protective levels of RPF2 was in the lower range of the recommended dosage for humans (pediatric −6 mg/kg, adult −20 mg/kg) (54, 55). Since TMP is an antibiotic, there might be concerns that chronic administration of the drug could alter the microbiota of the gut, adversely affecting the patient. Our animals on the TMP diet did not show any difference in feeding or behavior when compared to the control animals placed on normal chow (Supplemental Figure 6). Additionally, control animals fed with TMP had no changes to retinal function or morphology (Figure 3). Studies on the long-term use of TMP in patients with pneumocystis pneumonia, urinary tract infections, and acute lymphoblastic leukemia all found TMP to be safe, with minimal adverse effects (56–58). Moreover, to avoid systemic delivery, TMP could potentially be formulated as an eye drop for localized administration. Withdrawal of TMP reduced RPF2 levels to baseline, which further demonstrates the tightly-controlled regulation of this system. This is appealing when applying our therapy to a clinical setting because in the event of adverse effects the therapy has the ability to be stopped quickly.

Recently, the gene therapy field has achieved significant breakthroughs, including the FDA approval of the cell-based gene therapy known as Kymriah to treat precursor B-cell lymphoblastic leukemia (59). In addition, the first gene replacement therapy was granted FDA approval for treatment of an inherited retinal disorder (60). This is especially significant since it utilizes the AAV platform to deliver a normal copy of the *RPE65* gene to patients with Leber’s congenital amaurosis, who have a mutation in this gene. The success of this therapy opens doors for developing gene-based treatments for a variety of other retinal disorders. Our results suggest that broad-spectrum protection could be provided by RPF2 as a primary, standalone treatment to prevent retinal degeneration. Alternatively, it could be used as an adjunct to a gene replacement therapy. For example, in patients given AAV-*RPE65*, there was successful gene replacement and rescue of function, but cell loss persisted over time (61, 62). AAV-RPF2 could potentially be used to expand the therapeutic window for patients with inherited retinal disorders, keeping the cells alive long enough for secondary intervention (63). In addition, stem cells poorly survive in a degenerating environment (64). Long-term studies have shown that after integration, stem cells die over time (65), so a combinatorial approach using AAV-RPF2 could improve the efficacy of this treatment. Overall, this work is a promising step towards improved, broad-spectrum treatment options for patients suffering from retinal disease.

## Methods

### Mice

All procedures were performed according to the Association for Research in Vision and Ophthalmology Statement for the Use of Animals in Ophthalmic and Vision Research and in compliance with protocols approved by the University of Florida. BALB/cJ, C57BL/6, and *rd10* breeders were obtained from The Jackson Laboratory (Bar Harbor, ME). Colonies were established at the internal rodent breeding program at the University of Florida. All mice were reared in 12-hour cyclic dim light (<100 lux) / dark conditions (6AM-6PM) in the University of Florida animal housing facility and provided water and food *ad libitum*.

### AAV vector construction and virus production

The AAV vector plasmid ‘pTR-UF11’ contains the full chimeric CBA driving green fluorescent protein (GFP) cDNA (66). The AAV vector plasmid ‘pTR-CBA-hLIF’ contains the CBA promoter driving hLIF cDNA followed by an internal ribosome entry site and a GFP cDNA. This plasmid was generously provided by Dr. Sergei Zolotukhin (67). The AAV vector plasmid ‘pTR-CBA-RPF2’ contains the CBA promoter driving the RPF2 cDNA. The RPF2 cDNA was created by *de-novo* synthesis from Genscript (Piscataway, NJ). The rep-cap plasmid containing AAV2 cap with Y272F, Y444F, Y500F, Y730F and T491V mutations has been previously described (36). The vector plasmid, AAV rep-cap and Ad helper plasmid were co-transfected into HEK293T cells and the viruses were harvested and purified as previously described (68). Purified viruses were resuspended in balanced salt solution (BSS) (Alcon, Ft. Worth TX) supplemented with Tween-20 (0.014 %) and stored at −80°C until use. All viruses were titered by qPCR and tested for endotoxin.

### Cell culture transfections, TMP treatment, and conditioned media experiments

The rat müller glial-like cell line (rMC-1) was generously provided by Dr. Vijay Sarthy (69). The cells were cultured in high glucose Dulbecco’s minimal essential medium (DMEM) supplemented with 10% fetal bovine serum, 100 U/mL penicillin, 100 μg/mL streptomycin and 1x GlutaMax (Thermo Fisher Scientific Inc., Waltham, MA). At 80% confluency, the cells were transfected with a purified plasmid (Supplemental Table 1) using the standard Lipofectamine 3000 protocol (Thermo Fisher Scientific Inc., Waltham, MA). 24 hours after transfection, the media was replaced with serum-free DMEM. The cells were then dosed overnight with varying concentrations of TMP or control vehicle (20% DMSO). The conditioned media was then collected for ELISA or STAT3 activation studies. For conditioned media experiments, rMC-1 cells were grown in 6-well plates. The serum-deprived cells were incubated for 30 minutes with conditioned media collected from transfected rMC-1 cells treated with TMP. The cells were then washed with cold PBS and harvested for analysis by immunoblotting.

### Immunoblotting

Retinal tissue or rMC-1 cells were homogenized in RIPA buffer (Sigma-Aldrich, St. Louis, MO) containing a protease inhibitor cocktail (Merck Millipore, Billerica, MA) supplemented with sodium fluoride and sodium orthovanadate (New England BioLabs Inc., Ipswich, MA). Protein lysate concentration was quantified by BCA assay (Thermo Fisher Scientific Inc., Waltham, MA) before boiling and subjecting to SDS-PAGE. Lysates were run on 4%-20% polyacrylamide gel (Thermo Fisher Scientific Inc., Waltham, MA) and transferred onto polyvinylidene fluoride membranes (Merck Millipore, Billerica, MA). Membranes were blocked with Odyssey blocking buffer (LI-COR Biosciences, Lincoln, NE) and incubated overnight with primary antibodies (Supplemental Table 2). Secondary antibodies conjugated with IRDyes (Supplemental Table 2) were used. Signals were detected using the Odyssey CLx Imaging System (LI-COR Biosciences, Lincoln, NE). Intensities of the protein bands were quantified using Image Studio 5 (LI-COR Biosciences, Lincoln, NE).

### Enzyme-linked immunosorbent assay (ELISA)

LIF and RPF2 levels were detected in retinal tissue lysates and rMC-1 conditioned media. For tissue lysates, the retinas were prepared in an ELISA lysis buffer (1% IGEPAL- CA 630, 135mM NaCl, 20mM Tris and 2mM EDTA). Lysates or conditioned media was incubated for 2 hours on plates containing pre-blocked capture antibodies followed by 1-hour incubations with biotinylated detection antibody and streptavidin conjugated to horseradish peroxidase (Supplemental Table 2). Signals were developed using the Substrate Reagent Pack (R&D Systems, Inc., Minneapolis, MN) and read at a 450nm wavelength using the CLARIOstar microplate reader (BMG LABTECH GmbH, Ortenberg, Germany).

### TMP delivery

For the *in vivo* dose response, varying concentrations of TMP were resuspended in a solution of 20% DMSO. The mice were IP-injected once per day for seven days with equal volumes of the varying TMP dosage or vehicle. For the reversal study, mice were injected once per day for seven days followed by no administration of the drug for a subsequent seven days before the retinas were harvested for ELISA analysis. For the TMP food supplement, 600 ppm of TMP was added to the Teklad LM-485 mouse diet (Envigo, Huntingdon, United Kingdom). The TMP chow was provided *ad libitum* to the mice.

### Intravitreal injections

Mice were anesthetized with ketamine (70-100mg/kg) and xylazine (5-10mg/kg). The eyes were dilated with a single drop of tropicamide and phenylephrine (Alcon, Ft. Worth TX). 1µl of viral vector or BSS (Alcon, Ft. Worth TX) were separately injected into the vitreous through the temporal limbus of the eye using a syringe with a 33-gauge needle (Hamilton Company, Reno, NV) AK-Poly-Bac ophthalmic medication (Alcon, Ft. Worth TX) was used to prevent post-surgical infections and suppress inflammation. Eyes were evaluated by OCT one week post-injection, and any animals with unresolved surgical trauma were excluded from the study.

### Electroretinography (ERG)

Mice were anesthetized with ketamine (70-100mg/kg) and xylazine (5-10mg/kg). The eyes were dilated with a single drop of tropicamide and phenylephrine (Alcon, Ft. Worth TX). Retinal function was measured using the Colordome ERG system (Diagnosys, Dorset, United Kingdom). To record electrical responses from both eyes, gold wire electrodes were placed on the corneas, a platinum reference electrode was placed in the mouth and a platinum ground electrode was attached to the tail. The body temperature was maintained at 37°C throughout the experiment using the built-in heating pad of the ERG system. For scotopic ERGs, the animals were dark-adapted for at least 12 hours. A series of increasing light flash intensities ranging from 0.0001 - 180 cd.s/ m^2^ was used. The a-wave was defined as the trough of the negative deflection from the baseline, while the b-wave was measured from the a-wave to the peak of the positive deflection. For photopic ERGs, a background illumination of 30 cd.s/m^2^ was used to block rod photoreceptor contribution to the recording. A series of increasing light flash intensities above the background illumination (photopic 1.875 - 60 cd.s/m^2^) were used to generate a cone response.

### Spectral Domain-OCT and scoring

For *in vivo* retinal imaging, Spectral Domain-OCT images were obtained using the Envisu C-Class OCT system (Leica Microsystems, Wetzlar, Germany) Mice were first anesthetized with ketamine (70-100mg/kg) and xylazine (5-10mg/kg), followed by dilation with a single drop of tropicamide and phenylephrine (Alcon, Ft. Worth TX). Cornea clarity was maintained using GenTeal lubricating eye gel (Novartis Pharmaceuticals Corp., Basil, Switzerland). The mice were placed on a custom-built stage secured with a bite bar to allow free rotation for imaging. The stage was adjusted manually to center the image of the retina at the optic nerve head. Cross-sectional images were generated using 425 rectangular volume scans, which were averaged every five scans prior to image analysis. Images were acquired for both eyes. Averaged OCT images were analyzed using InVivoVue 2.4 (Leica Microsystems, Wetzlar, Germany). Outer nuclear layer (ONL) thickness was measured using the linear caliper function in the software by a masked observer using a pre-established uniform grid (Supplemental Figure 2 Top panel). The OCT images were also graded by a masked observer utilizing a scoring system ranging from 1 to 3 (Figure 1C). A score of 3 indicated a healthy-normal retina with no thinning and little to no inflammation. A scoring of 2 indicated a retina with slight edema, minor thinning, or inflammation. A scoring of 1 signified a degenerated retina with severe thinning, severe inflammation, missing external limiting membrane, or visible retinal detachment.

### Immunohistochemistry (IHC)

To obtain retinal tissue sections, mouse eyes were enucleated with forceps and placed in cold PBS. Under a dissecting microscope, the cornea and lens were dissected away and the remaining eye cup was processed for cryosectioning. In additional eyes, retinal flat-mounts were prepared by gently dissecting the retina from the eyecup. Four incisions were made from the periphery to the center of the retina to flatten the retina onto a microscope slide. Eyecups and flat mounts were fixed in 4% paraformaldehyde for 30 minutes at room temperature. The fixed eyecups were cryopreserved in 30% sucrose in PBS overnight at 4°C. The samples were embedded in Tissue-Tek OCT Compound (Sakura Fintek USA Inc., Torrance, CA) and immediately frozen in liquid nitrogen. Tissue sections were cut at a thickness of 16μm on a cryostat and placed onto glass slides. Slides were stored at −20°C until needed for immunostaining. For IHC, the flat mount retinas or eyecup sectioned slides were washed with PBS containing 1% Triton X-100 (PBS-T). To prevent non-specific binding, the tissue was blocked with 10% horse serum in PBS-T for at least 1 hour at room temperature. Primary antibody (Supplemental Table 2) was applied to each sample for incubation overnight at 4°C. Samples were washed with PBS-T before incubating with secondary antibody (Supplemental Table 2) for 1 hour at room temperature. Nuclei were counter-stained with 4-6-Diamidino-2-phenylindole (DAPI) before mounting with glass coverslips using 60% glycerol. Imaging was performed using the BZ-9000 fluorescence microscope (Keyence Corp., Osaka, Japan). The exposure time of each fluorescent channel was kept constant between samples for a given antibody. For flat mount counting of cone outer segments, a 250 mm^2^ grid was superimposed over five different areas of each flat mount image. Cells within the grid were counted. All data collection was done by a masked observer.

### Light damage

For light damage experiments, two mice were placed in a cage with a modified lid equipped with LED lights (white light). Prior to damage, the light intensity was measured using a light meter (Thermo Fisher Scientific Inc., Waltham, MA) and set to 1200 lux using the connected dimmer. The mice were exposed to this damaging light for 4 hours from 6PM-10PM. Once the experiment ended, the mice were moved back into dim lighting conditions to recover for one week before downstream analysis was performed. The lighting equipment is available at the UF animal facility and approved by the ACS. The mice were kept in a ventilated rack for the duration of the experiment.

### OptoMotry

To test the optokinetic reflex of mice we used the OptoMotry system developed by CerebralMechanics Inc. (Lethbridge, Canada). The system consisted of four computer screens that display alternating vertical light and dark bar that appear as a virtual, rotating, 3D cylinder. The mouse was placed on a non-rotating pedestal in the middle of the enclosure. The 3D cylinder pattern was rotated at a fixed speed of 12 degrees per second and the contrast was maintained at 100%. Head tracking was monitored using an overhead camera, which was recorded by a masked observer. The staircase method was used to determine visual acuity by establishing the threshold of spatial frequency that the animal can track. An animal was considered to be tracking if its head followed the direction of the stimulus with a speed similar to the pattern movement.

### Fundus imaging

For fundus imaging, a Micron III camera (Phoenix Research Labs, Pleasanton, CA) was used. Mice were anesthetized with ketamine (70-100mg/kg) and xylazine (5-10mg/kg) and dilated with a single drop of tropicamide and phenylephrine (Alcon, Ft. Worth TX). Corneal clarity was maintained using GenTeal lubricating eye gel (Novartis Pharmaceuticals Corp., Basil, Switzerland). The mice were stabilized on a custom holder that allowed adjustments to center the image of the retina at the optic nerve head. Imaging was performed in under brightfield illumination and using filtered red and green channels. Images were acquired for both eyes.

### Statistical analysis

Statistical analysis was carried out using Prism 6 (GraphPad Software Inc., La Jolla, CA). Statistical significance between multiple groups was determined by using Analysis of Variance (ANOVA).

## Acknowledgements

We would like to thank the Dr. William Hauswrith and the University of Florida’s Ocular Gene Therapy Core for manufacturing the AAV vectors, Dr. Sergei Zolotukhin for the pTR-CBA-hLIF vector and Dr. Vijay Sarthy for providing the rMC-1 cell line used in this study. We also thank Dr. Wesley Clay Smith, Dr. Jorg Bungert, Dr. Alfred Lewin, Dr. Christhian Idelfonso, Dr. Shreyasi Choudhury, Dr. Seok Hong Min, Raghav Pai, Eric Nayman, Lei Xu, Marcus Hooper, Emily Brown and Megan Ash for their assistance on this project. This work was supported by the Foundation for Fighting Blindness, the National Eye Institute (Grant # R01EY016459), and an unrestricted departmental grant from the Research to Preventing Blindness. The work was performed at the University of Florida in Gainesville, FL.

## Author Contributions

C.P.S. and J.D.A. conceived RPF2 and designed the experiments. C.P.S., C.J.K. and A.A.I. performed experiments. C.P.S., C.J.K., and J.D.A. prepared the manuscript. S.L.B contributed advice on AAV serotype and S.L.B and S.E.B provided feedback on the manuscript. J.D.A acquired funding.

