## Supplementary Materials for "A drug-tunable gene therapy for broad-spectrum protection against retinal degeneration" (from another version of this preprint: version 1, 2018-03-16)

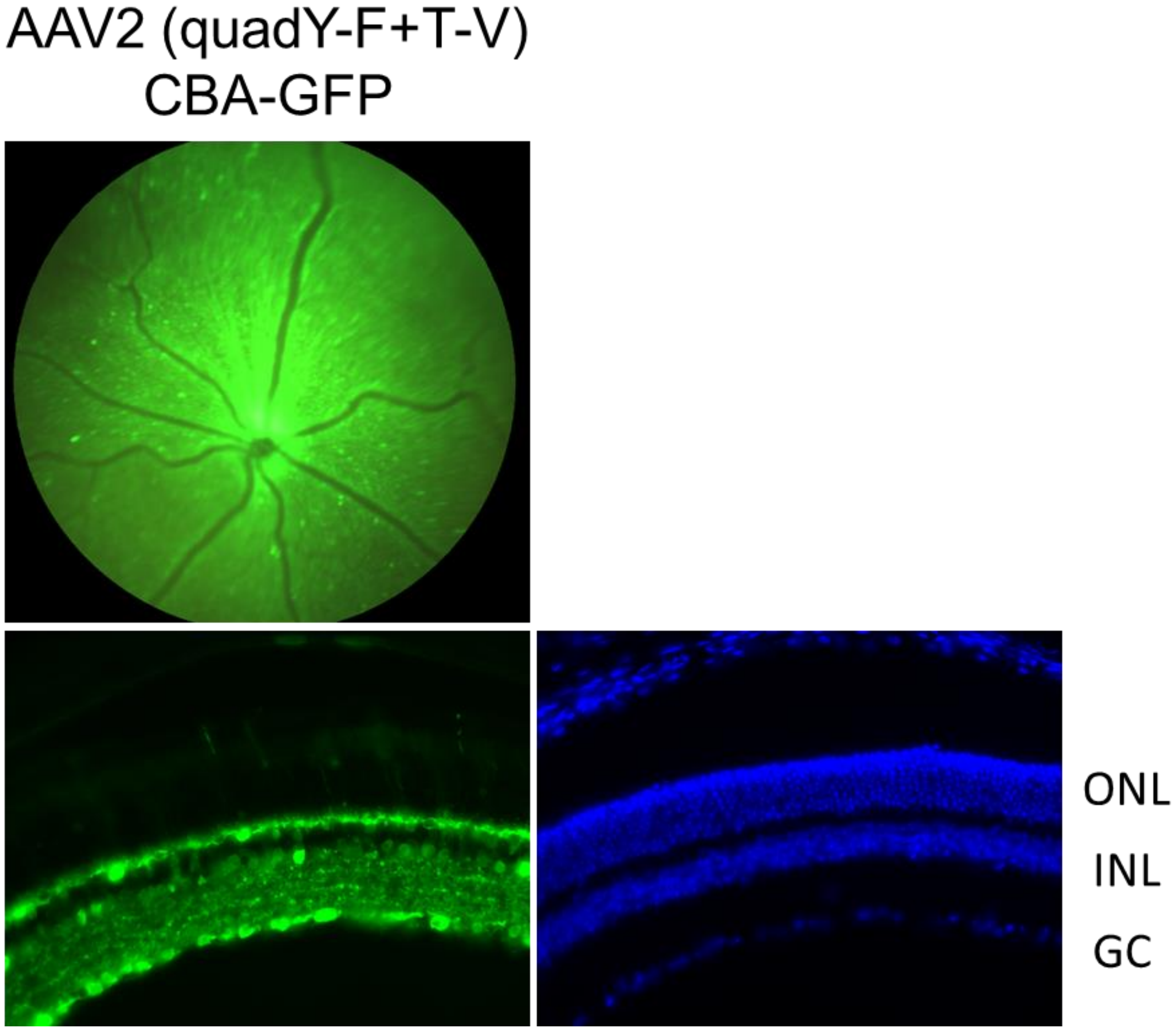


**Supplemental Figure 1. AAV2 (quadY-F+T-V) tropism in the mouse retina.** Top: Fundus images of the retina injected intravitreally with AAV2 (quadY-F+T-V) CBA-GFP showing. Bottom: Immunohistochemisty cross section showing GFP expression (green) and DAPI stained nuclei (blue). ONL = Outer nuclear layer, INL = Inner nuclear layer and GC = Ganglion cell layer. Images were obtained at 40X magnification.

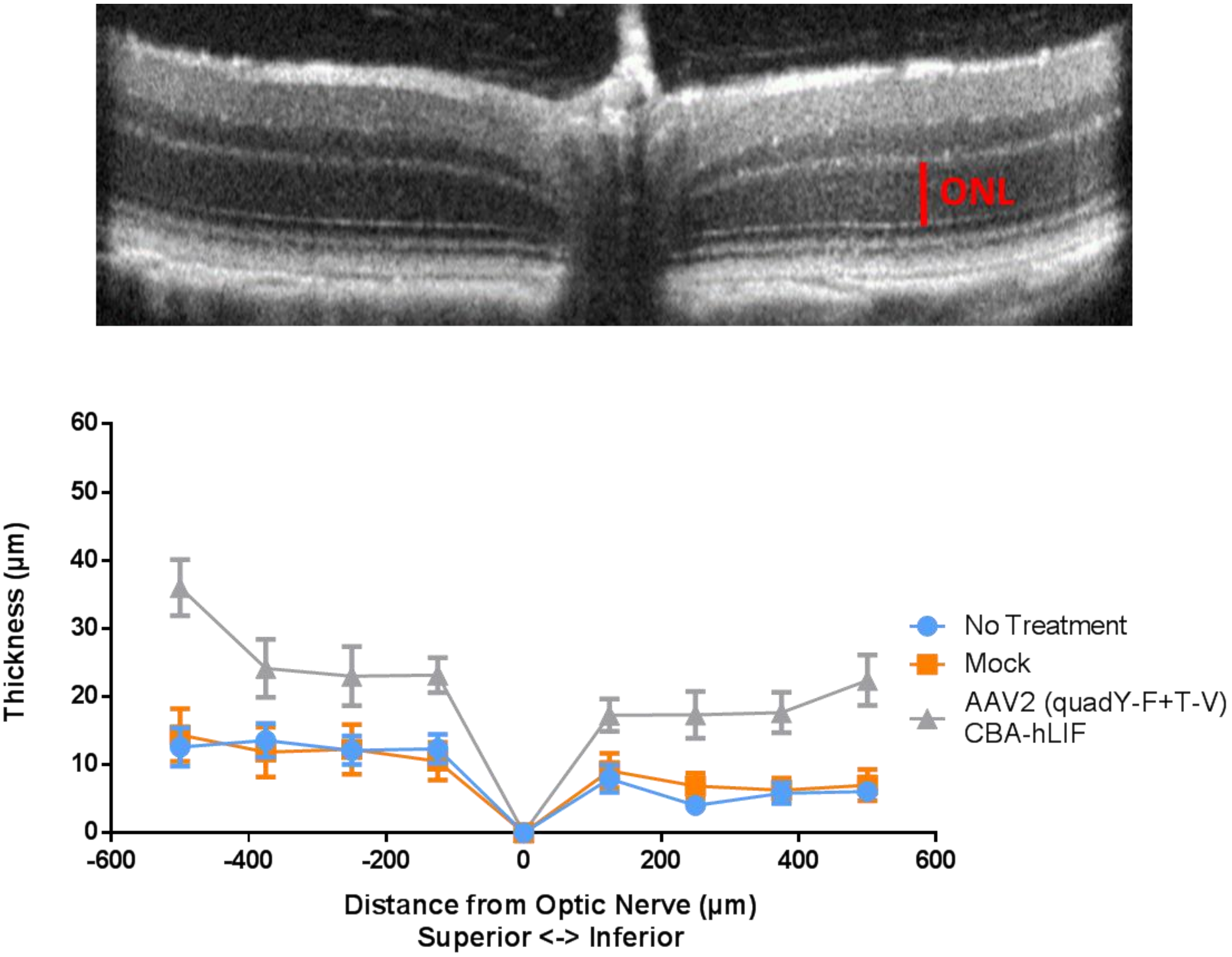


**Supplemental Figure 2. AAV delivery of hLIF protects the retina from light damage.** Top: Representative OCT image showing how the outer nuclear layer (ONL) thickness is measured. Thickness was measured from superior to inferior retina 500μm from the optic nerve. Bottom: Spider plot of the ONL thickness of light damaged retinas treated with AAV2 (quadY-F+T-V) CBA-hLIF. No treatment and mock injected were used as controls.

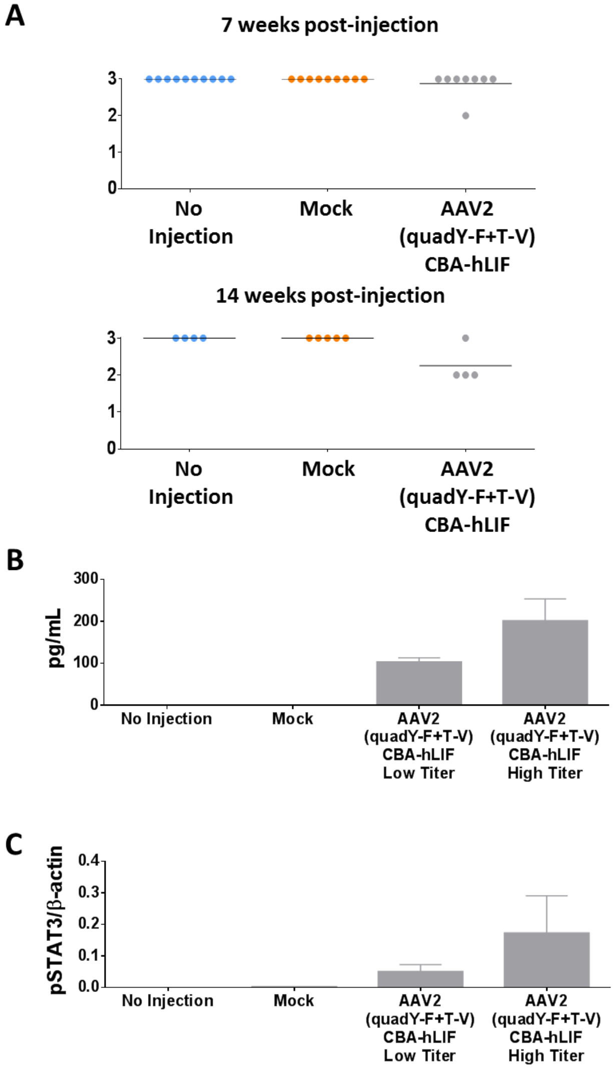


**Supplemental Figure 3. Characterization of AAV2 (quadY-F+T-V) CBA-hLIF delivery in the mouse retina.** **A)** Scoring of retinas with high titer AAV2 (quadY-F+T-V) CBA-hLIF. **B)** Levels of hLIF in the retina determined by ELISA. **C)** Activation of STAT3 as determined by immunoblot quantification.

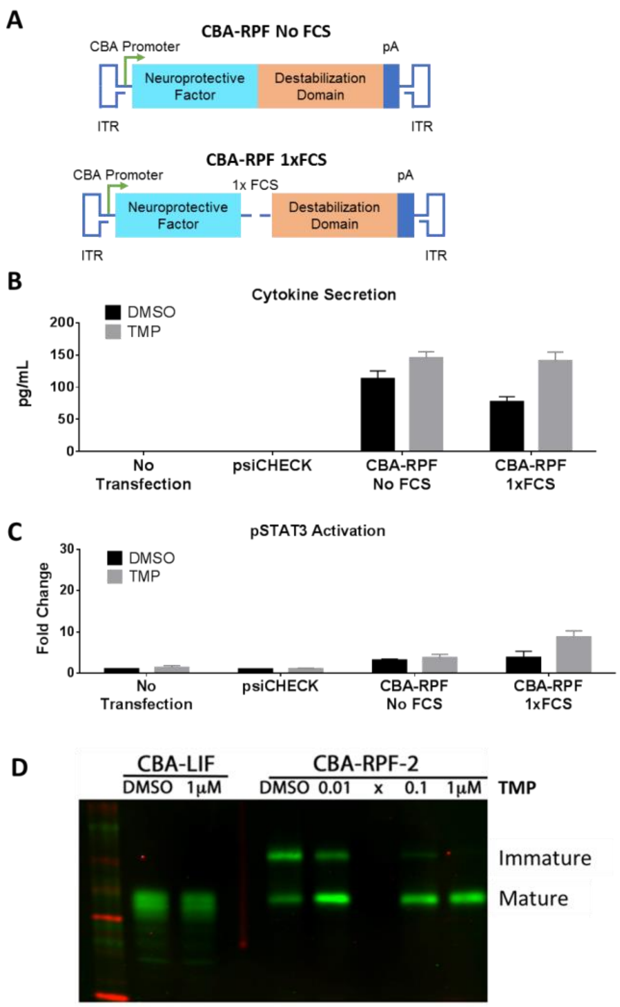


**Supplemental Figure 4. Iterations of RPF2 development. A)** Schematics of CBA-RPF iterations with no or one furin cleavage site (FCS). Both vectors are approximately 3.4kb in size **B)** Levels of cytokine secreted by rMC-1 transfected cells into the conditioned media detected by ELISA. **C)** Activation of STAT3 in rMC-1 cells treated with conditioned media. **D)** Dose-dependent stabilization and maturation of RPF2 by TMP in transfected rMC-1 cells.

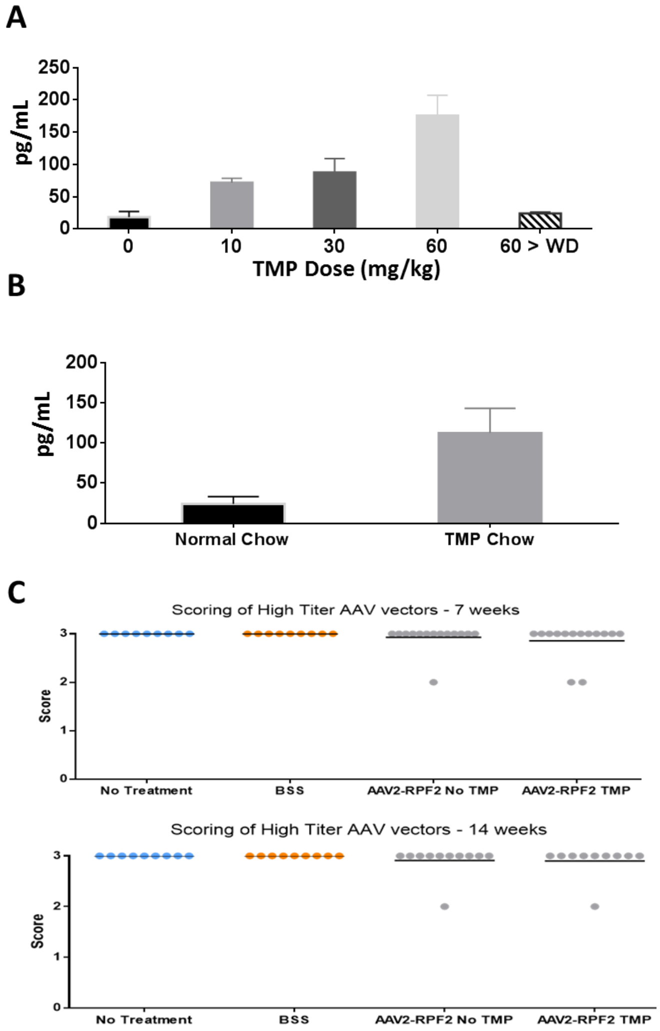


**Supplemental Figure 5. Characterization of RPF2 *in vivo.* A)** Levels of RPF2 in the retina of animals injected with the high titer AAV2 (quadY-F+T-V) CBA-RPF2 dose. **B)** Levels of RPF2 in the retina when TMP is administered in the food. Cytokine levels were detected by ELISA after one week of TMP treatment. **C)** Scoring of retinas inject with high titer of AAV2 (quadY-F+T-V) CBA-RPF2.

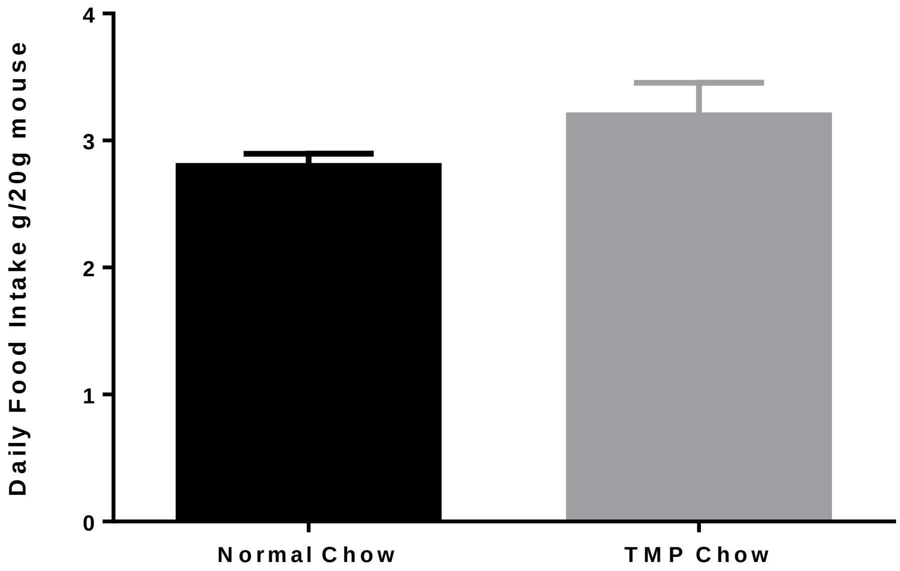


**Supplemental Figure 6. Daily food intake is unchanged with TMP supplemented chow.** Food intake was measured for one week for mice on normal and TMP chow.

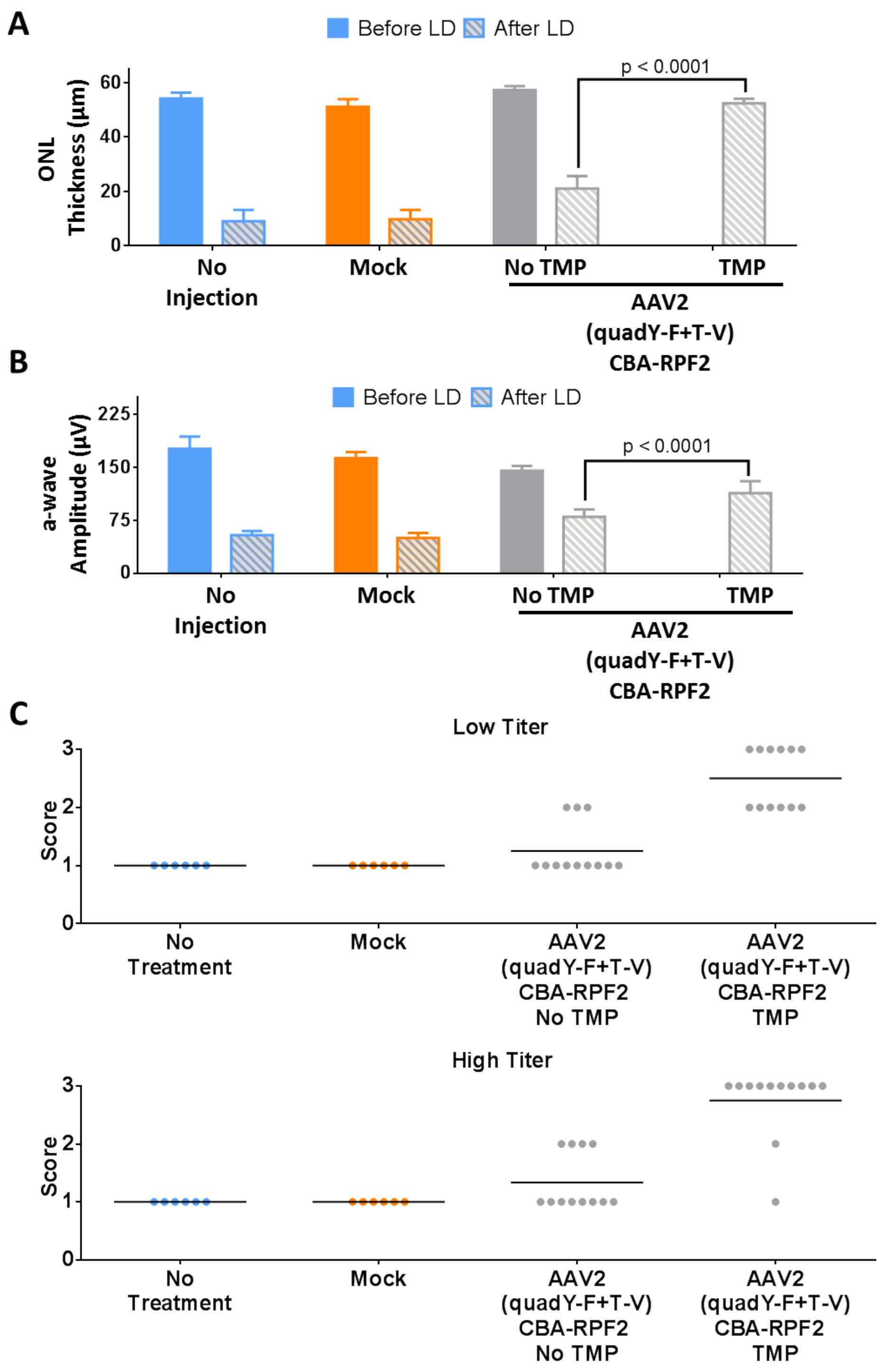


**Supplemental Figure 7. RPF2 treated retinas show improved retinal structure and function after bright light exposure. A)** Averaged thickness of the retinal outer nuclear layer and **B)** retinal function by ERG after RPF2 stabilization in high titer treated animals. **C)** Low titer and High titer RPF2-treated animals showed less signs of edema, rosette formation and infiltrating immune cells compared to the control groups.

| Plasmid Name | Promoter | Protein expressed | Use |
|---|---|---|---|
| pTR-UF11 | chicken beta-actin | Green Fluorescent Protein | Animals |
| pTR-CBA-hLIF | chicken beta-actin | human Leukemia Inhibitory Factor | Cell culture/Animals |
| pTR-CBA-RPF2 | chicken beta-actin | Retinal Protective Factor 2 | Cell culture/Animals |
| psiCHECK (Promega Corporation) | SV40 early enhancer/promoter | Renilla luciferase | Cell culture |

**Supplemental Table 1: Plasmids used in this study**

| Name | Company | Catalog No. | Species raised | Use |
|---|---|---|---|---|
| Biotin hLIF | R&D Systems | BAF250 | Goat | 3µg/ml Detection for ELISA/1:500 dilution Immunoblots |
| hLIF | R&D Systems | AF-250-NA | Goat | 3µg/ml Capture for ELISA |
| Streptavidin-HRP | R&D Systems | DY998 | | 1:200 dilution Detection signal for ELISA |
| pSTAT3 | Cell Signaling | 9145L | Rabbit | 1:2,000 dilution Immunoblots |
| beta actin | Abcam | ab6276 | Mouse | 1:10,000 dilution Immunoblots |
| Streptavidin-800CW | Li-Cor | 926-32230 | | 1:5,000 dilution Immunoblots |
| anti-mouse IRDye 680RD | Li-Cor | 926-68170 | Goat | 1:10,000 dilution Immunoblots |
| anti-rabbit IRDye 800CW | Li-Cor | 925-32211 | Goat | 1:10,000 dilution Immunoblots |
| s-opsin | Santa Cruz | sc-14363 | Goat | 1:300 dilution Immunohistochemistry |
| m-opsin | EMD Millipore | AB5405 | Rabbit | 1:300 dilution Immunohistochemistry |
| anti-goat Alexa Fluor 488 | Life Technologies | A21206 | Donkey | 1:500 dilution Immunohistochemistry |
| anti-rabbit Alexa Fluor 488 | Life Technologies | A11055 | Donkey | 1:500 dilution Immunohistochemistry |

**Supplemental Table 2: Antibodies used in this study**
